# Gait adaptation to asymmetric hip stiffness applied by a robotic exoskeleton

**DOI:** 10.1101/2023.10.10.561679

**Authors:** Banu Abdikadirova, Mark Price, Jonaz Moreno Jaramillo, Wouter Hoogkamer, Meghan E. Huber

## Abstract

Wearable exoskeletons show significant potential for improving gait impairments, such as interlimb asymmetry. However, a more profound understanding of whether exoskeletons are capable of eliciting neural adaptation is needed. This study aimed to characterize how individuals adapt to bilateral asymmetric joint stiffness applied by a hip exoskeleton, similar to split-belt treadmill training. Thirteen unimpaired individuals performed a walking trial on the treadmill while wearing the exoskeleton. The right side of the exoskeleton acted as a positive stiffness torsional spring, pulling the thigh towards the neutral standing position, while the left acted as a negative stiffness spring pulling the thigh away from the neutral standing position. The results showed that this intervention applied by a hip exoskeleton elicited adaptation in spatiotemporal and kinetic gait measures similar to split-belt treadmill training. These results demonstrate the potential of the proposed intervention for retraining symmetric gait.

## I. INTRODUCTION

Stroke is the primary cause of neurological disability among adults, leading to mobility limitations and negatively affecting their overall quality of life [1]. More than 80% of stroke survivors experience walking dysfunction, which causes difficulties in performing daily living activities [2]. Common gait abnormalities due to stroke include slower walking speed [3], as well as asymmetries in gait kinematic [4] and kinetic [5] behavior.

Asymmetric gait increases the risk of musculoskeletal injury in the non-paretic leg due to excessive joint loading [6] and can decrease the musculoskeletal health of the paretic leg due to disuse [7]. It is also known to be metabolically inefficient [8] and can negatively impact balance during walking [9]. Therefore, interventions that can effectively reduce asymmetries in gait patterns are greatly needed.

Split-belt treadmill training, which involves running the belts of a dual-belt treadmill at different speeds, has emerged as a promising strategy for restoring gait symmetry in individuals post-stroke [10], [11]. For individuals with symmetric gait, this paradigm initially induces step length asymmetry. With time, their gait gradually adapts to restore symmetry. Upon returning the belts to the same speed, a short-lived after-effect emerges, characterized by step length asymmetry in the opposite direction [12]. Subsequent studies found that split-belt treadmill training can elicit adaptation to decrease step length asymmetry in individuals post-stroke [10], [13]. Initially, when the paretic leg is placed on the slow belt, the individual takes longer steps with the paretic leg and shorter steps with the non-paretic leg. With time, the individual adapts their gait to the amplified asymmetry, resulting in after-effect of improved step length symmetry once the belts are returned to the same speed [14].

Despite the promise of split-belt treadmill training for restoring gait symmetry in individuals post-stroke, the transfer of these improvements observed in the after-effect to over-ground walking remains limited [15], [16]. Repetitive practice has been shown to enhance this transfer and lead to longer-term improvements [15]. However, the restricted accessibility of the specialized treadmill equipment needed to deliver split-belt training limits individuals’ ability to engage in repetitive practice.

Wearable robotic technology presents a potential solution to overcome the existing limitations associated with split-belt treadmill training. Compared to split-belt treadmills, wearable robotic exoskeletons are smaller in size, more affordable, and easier to use during activities of daily living. Exoskeletons can also be designed to be autonomous and portable, which would allow overground training outside of clinical or laboratory settings.

Recent studies have highlighted the potential of lower limb exoskeletons in compensating for asymmetries and improving walking performance in individuals post-stroke [17], [18]. However, the primary focus of compensation or assistance paradigms is to restore or attain a desired motor behavior while the wearable exoskeleton is actively being used. In the context of rehabilitation, the ultimate goal is to restore a desired motor behavior when the wearable exoskeleton is removed. It is still an open question of whether a wearable robotic exoskeleton can feasibly induce neural changes in the nervous system to improve symmetry in a persistent manner.

Rather than compensating for gait asymmetry, we propose that exaggerating gait asymmetry with an exoskeleton, similar to split-belt treadmills, may be able to recreate the adaptation responses leading to reduced gait asymmetry. To determine the feasibility of such an intervention, the goal of the present study was to investigate whether applying an asymmetric stiffness with a hip exoskeleton elicits locomotor adaptation and assess whether such adaptation is generally comparable to that observed during split-belt treadmill training in individuals with symmetric gait.

In the present study, the hip exoskeleton was controlled to emulate a positive, torsional spring to act as an attractor on the right hip joint and a negative, torsional spring to act as a repeller on the left hip joint. Emulating torsional springs at the hip joints promotes user comfort and safety, as it does not overly constrain or disrupt the natural dynamics of walking [19]. A stiffness-based intervention also encourages participants to actively engage in the task [20] and allows them to adjust their gait in response to the applied intervention.

We hypothesized that the initial application of these asymmetric stiffnesses about the hip joints would induce asymmetry in gait kinematics and kinetics similar to that induced by the split-belt treadmill paradigm (*Hypothesis 1*). We further predicted that with the perturbation applied, individuals would adapt their gait behavior back towards restoring gait symmetry over time (*Hypothesis 2*) and exhibit an aftereffect in an opposite direction to initially induced asymmetry after the perturbation is removed (*Hypothesis 3*). Preliminary reports of this study and an initial pilot study were presented in [21] and [22], respectively.

## II. EXPERIMENTAL METHODS

### A. Participants

Thirteen non-impaired individuals (sex: two female, eleven male; age: 23.1 *±*3.3 years; height: 1.72 *±*0.09 m; mass: 66.9 *±*15.3 kg) took part in this study. None had previously worn a hip exoskeleton or participated in a similar experiment. All participants signed an informed consent form before the experiment. The experimental protocol was reviewed and approved by the Institutional Review Board of the University of Massachusetts Amherst (Protocol ID: 3066, Approval Date: November 01, 2021).

### B. Robotic Hip Exoskeleton

The hip exoskeleton used in this study was a custom robot developed by the Human Robot Systems Laboratory (HRSL) at University of Massachusetts Amherst (Figure 1A). The exoskeleton (3kg) is worn around the waist and fastened to the thighs. The waist and thigh components are size-adjustable. Two actuators, one at each hip joint, can apply flexion and extension torque about the hip joints in the sagittal plane; passive hinges allow for hip adduction and abduction movements in the frontal plane. Each actuator consists of a brushless DC motor with a 6:1 gearhead and an absolute encoder, along with additional sensors and electronics (ActPack 4.1, Dephy, Maynard, MA, USA). Output torque from the actuator is determined and controlled by sensing the electrical current in the motor. High-level operation is controlled through a Raspberry Pi 4 (Raspberry Pi Ltd, Cambridge, UK) microcomputer. As this study was performed on a treadmill, the power source and the microcomputer were located offboard.

**Fig. 1.**
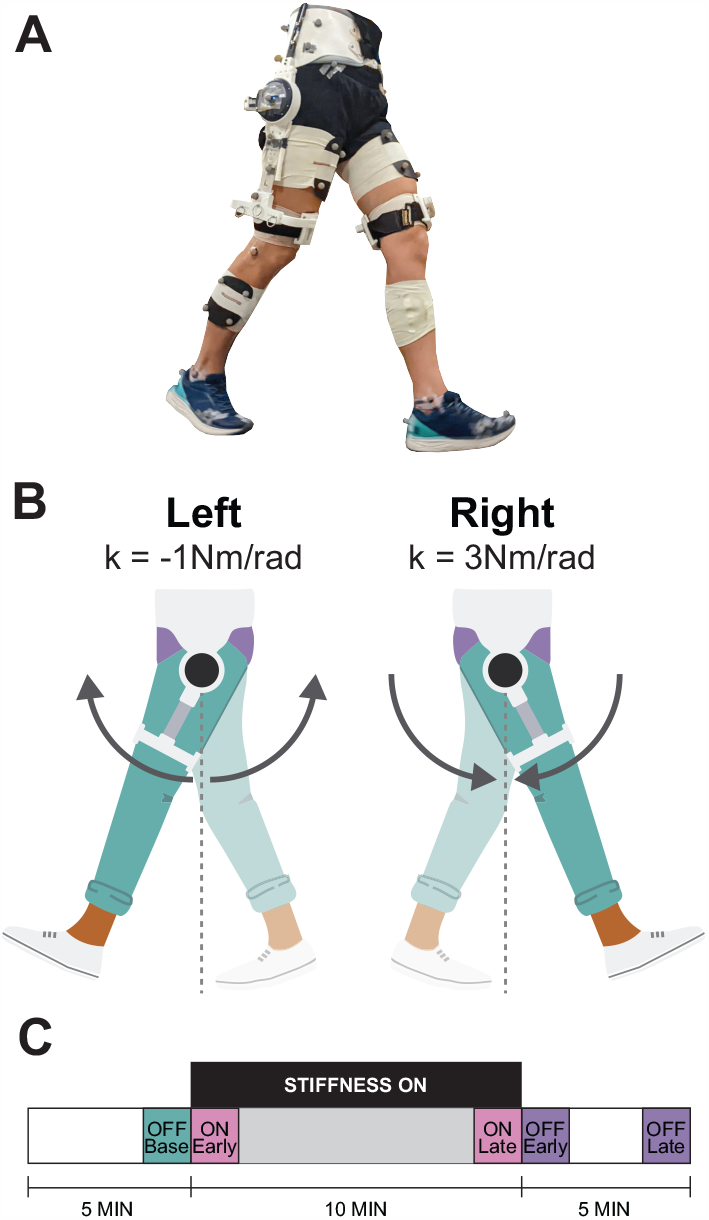
**A** Participants walked on an instrumented treadmill wearing a bilateral hip exoskeleton and reflective markers for motion capture. **B** The exoskeleton exerted attractive and repulsive torques expressed as positive and negative stiffness on the right and left sides, respectively. **C** Experimental protocol consisted of 5 minutes of baseline walking (stiffness controller OFF), 10 minutes of walking with stiffness ON, and 5 minutes of walking with stiffness OFF.

### C. Asymmetric Hip Stiffness Controller

The actuators on the hip exoskeleton were controlled to emulate virtual, torsional springs using the following control law:

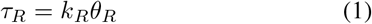

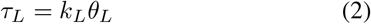

where *τ*_*R*_ and *τ*_*L*_ are the torques applied by right and left actuators, *θ*_*R*_ and *θ*_*L*_ are the right and left hip angles relative to an upright standing position, and *k*_*R*_ and *k*_*L*_ are the stiffness values of the virtual springs about the right and left hip joints, respectively.

In this study, the stiffness values were set to *k*_*R*_ = 3 Nm/rad and *k*_*L*_ = -1 Nm/rad. The positive, virtual spring on the right hip acted to pull the right thigh towards the upright standing position, whereas the negative, virtual spring on the left acted to push the left thigh away from the upright standing position (Figure 1B). Increasing hip stiffness restricts joint motion, whereas decreasing hip stiffness amplifies it [23]. Thus, the positive and negative springs were intended to mimic the fast (which initially shorten step lengths) and slow belts (which initially lengthen step lengths), respectively, in the split-belt treadmill training paradigm. The specific stiffness values used in this experiment were chosen based on pilot testing to ensure safety and user comfort [22]. The stiffness controller was turned off by setting *τ*_*R*_ = *τ*_*L*_ = 0.

### D. Experimental Protocol

Participants walked at 1.3m/s on an instrumented dual-belt treadmill (Bertec Corporation, Columbus, OH, USA) in the following conditions. Participants first walked for two minutes without wearing the exoskeleton to acclimate to walking on the dual-belt treadmill, after which they were fitted with the exoskeleton. Participants then walked for 20 minutes wearing the exoskeleton, during which the stiffness controller was off for the first five minutes, on for the next ten minutes, and off again during the remaining five minutes (Figure 1C).

#### 1) Kinematic Data

Fifty-two reflective markers were placed on the lower limbs of the participant and the exoskeleton to locate the pelvis, thighs, shanks, feet, and exoskeleton segments in 3D space. Marker data was recorded at 100Hz with an eight-camera motion capture system (Qualisys, Inc., Gothenburg, Sweden).

Recorded marker data was labelled and processed in the Qualisys software and then filtered with a fourth-order zero-lag Butterworth low-pass filter (6 Hz) using the filtfilt function in MATLAB (The Mathworks, Natick, MA) to remove high frequency noise.

A 3D model of the human-exoskeleton system was created in OpenSim 4.3 [24] for each participant based on the “Gait2354” model to calculate lower limb joint angles via inverse kinematics. The exoskeleton is composed of two three-segment systems, which include a waist harness, a motor segment, and a thigh frame. These segments are linked by two revolute joints: an ab/adduction hinge connecting the motor to the waist harness and a flexion/extension joint between the motor and thigh frame, which represents the motor angle. The exoskeleton components are either constrained relative to the human skeleton or to each other. Each waist harness segment is connected to the pelvis by a six-degree-of-freedom joint, while the thigh segment is completely disconnected from the human thigh segment. This arrangement enables the exoskeleton and human joint kinematics to be evaluated independently. The OpenSim model and sample marker-set are available for download at: https://simtk.org/projects/gait-hip-exo.

Marker data recorded during standing was used to scale the OpenSim model and adjust the markers for each participant. Inverse kinematics was performed using a global least-squares optimization in OpenSim [25]. This approach involved minimizing the differences between model marker positions and those observed in the experiment, while taking into account joint constraints imposed by the model. To prevent unrealistic metatarsophalangeal (MTP) joint movement due to missing foot markers in some participants, MTP joint angle was constrained to be zero. This ensured accurate ankle joint kinematics, despite the limitations caused by the missing markers.

#### 2) Kinetic Data

Ground reaction forces (GRFs) were recorded at 1000Hz from two force plates located under the treadmill belts (Bertec Corporation, Columbus, OH, USA). Ground rection force data was filtered with a fourth-order zero-lag Butterworth low-pass filter (30 Hz) using the filtfilt function in MATLAB (The Mathworks, Natick, MA) to remove high frequency noise.

### E. Dependent Measures

Gait kinematic and kinetic measures for each leg were computed for every stride of each participant. Furthermore, asymmetry was computed to compare the kinematic and kinetic measures between the right and left leg for each stride.

#### 1) Stride Segmentation

One stride (i.e., one gait cycle) was defined as starting at the right leg heel-strike (0%) and ending at the subsequent heel-strike of the same leg (100%), determined using a threshold of 10 N on the rising edge of the right belt vertical GRF.

#### 2) Kinematic measures

Step length for each leg was quantified by the anterior-posterior distance between the heel markers at heel-strike of the given leg. Step time for each leg was quantified by time interval between heel-strike of the given leg and the subsequent heel-strike of the opposite leg. While this definition of step time may appear counter-intuitive, it was chosen based on the prevailing convention in the field which also makes it easier to compare to prior studies. Range of motion (ROM) of the hip, knee, and ankle joints of each leg was quantified by the difference between the maximum and minimum angular positions in the sagittal plane for each stride. The hip angle was defined as the relative angle between the femur and the pelvis, knee angle was defined between the tibia and the femur, and the ankle joint was defined between the foot and the tibia.

#### 3) Kinetic measures

The load-bearing properties of each stride were quantified by analyzing the peak propulsive, braking, and vertical GRF for each limb, which were normalized according to the participant’s body mass. Peak propulsive and braking ground reaction forces were extracted from anterior-posterior force measures. Peak propulsion and braking GRFs corresponded to the positive and negative peak of filtered anterior-posterior GRFs, respectively. Peak vertical GRF was determined to be the maximum value of filtered vertical force data per stride. The kinetic data of one participant was removed from the analysis due to frequent crossover of the belts, which rendered the individual ground reaction force contributions from the left and right legs indistinguishable.

#### 4) Asymmetry in the dependent measures

For each afore-mentioned kinematic and kinetic measure, the asymmetry between the positive stiffness and negative stiffness legs in each stride was quantified by the following ratio

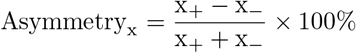

where x_+_ and x_−_ represent the measure for the positive stiffness (right) and negative stiffness (left) leg, respectively. A positive asymmetry in the measure indicates that the magnitude of the measure was higher for the leg with the positive stiffness applied. The opposite is true for a negative asymmetry.

### F) Statistical Analysis

#### 1) Conditions

The average asymmetry of each dependent measure was computed for every participant at five different time intervals. These time intervals, referred to as conditions, include

- *OFF:Base*: the final 10 strides during the baseline period with the stiffness controller switched off,
- *ON:Early*: the first 10 strides during the exposure phase with the stiffness controller switched on,
- *ON:Late*: the last 10 strides during the exposure phase with the stiffness controller switched on,
- *OFF:Early*: the first 10 strides during the post-exposure phase with the stiffness controller switched off
- *OFF:Late*: the last 10 strides during the post-exposure phase with the stiffness controller switched off

A summary of these conditions is illustrated in Figure 1C.

#### 2) Analyses of variance (ANOVAs)

A one-way repeated measures (ANOVA) was carried out to evaluate the effect of condition on the asymmetry in each dependent measure. If the results of the Mauchly’s test indicated that the assumption of sphericity (i.e., the assumption of equal variances of the differences between all combinations of conditions) was violated, then the Greenhouse-Geisser correction factor was applied to the degrees of freedom of the ANOVA.

#### 3) Planned comparisons

Upon detecting a significant effect of the condition, planned comparisons using paired *t*-tests were performed to further evaluate our three hypotheses. For all measures, except knee and ankle ROM, one-tailed tests were used since the direction of changes in asymmetry between conditions could be predicted. Predictions were either based on previous studies on split-belt treadmill walking (e.g., step length [26], step time [27], and ground reaction forces [26]–[28]) or the effect of applied hip stiffness during walking (hip ROM [23]) as summarized in Table II. Two-tailed tests were used for knee and ankle ROM measures due to the absence of prior results, which prevented a priori predictions of the direction of change in asymmetry.

**TABLE I.** Summary of ANOVA Results.

| Measure | Source | df | F | p |
| --- | --- | --- | --- | --- |
| Step Length Asymmetry | Condition | 2.34 | 46.16 | <0.001* |
|  | Error | 28.03 |  |  |
| Step Time Asymmetry | Condition | 4 | 20.44 | <0.001* |
|  | Error | 48 |  |  |
| Hip RoM Asymmetry | Condition | 1.42 | 41.60 | <0.001* |
|  | Error | 17.08 |  |  |
| Knee ROM Asymmetry | Condition | 4 | 41.00 | <0.001* |
|  | Error | 48 |  |  |
| Ankle ROM Asymmetry | Condition | 2.23 | 0.98 | 0.40 |
|  | Error | 26.81 |  |  |
| vGRF Asymmetry | Condition | 1.59 | 22.11 | <0.001* |
|  | Error | 17.45 |  |  |
| Braking Asymmetry | Condition | 4 | 49.26 | <0.001* |
|  | Error | 44 |  |  |
| Propulsion Asymmetry | Condition | 4 | 10.78 | <0.001* |
|  | Error | 44 |  |  |

\* indicates statistical significance

**TABLE II.** Planned Comparison Results.

|  | Test | Prediction | M | SD | df | t | p-adjusted |
| --- | --- | --- | --- | --- | --- | --- | --- |
| Step Length Asymmetry | H1 | one-tail (↓) | -3.61% | 1.80% | 12 | -7.24 | <0.001* |
|  | H2 | one-tail (↑) | 1.74% | 2.35% | 12 | 2.68 | 0.040* |
|  | H3a | one-tail (↑) | 5.43% | 2.77% | 12 | 7.07 | <0.001* |
|  | H3b | one-tail (↑) | 3.56% | 1.83% | 12 | 7.02 | <0.001* |
| Step Time Asymmetry | H1 | one-tail (↑) | 1.31% | 0.98% | 12 | 4.82 | <0.001* |
|  | H2 | one-tail (↓) | -0.41% | 0.83% | 12 | -1.78 | 0.200 |
|  | H3a | one-tail (↓) | -1.90% | 0.90% | 12 | -7.63 | <0.001* |
|  | H3b | one-tail (↓) | 1.00% | 1.08% | 12 | -3.33 | 0.012* |
| Hip RoM Asymmetry | H1 | one-tail (↓) | -11.75% | 6.51% | 12 | -6.51 | <0.001* |
|  | H2 | one-tail (↑) | 2.70% | 3.36% | 12 | 2.90 | 0.0268* |
|  | H3a | one-tail (↑) | 11.73% | 6.44% | 12 | 6.56 | <0.001* |
|  | H3b | one-tail (↑) | 2.68% | 1.87% | 12 | 5.19 | <0.001* |
| Knee ROM Asymmetry | H1 | two-tail | 4.46% | 2.47% | 12 | 6.50 | <0.001* |
|  | H2 | two-tail | -2.22% | 2.26% | 12 | -3.53 | 0.017* |
|  | H3a | two-tail | -5.25% | 2.36% | 12 | -8.04 | <0.001* |
|  | H3b | two-tail | -3.01% | 1.78% | 12 | -6.08 | <0.001* |
| vGRF Asymmetry | H1 | one-tail (↓) | -4.15% | 4.12% | 11 | -3.49 | 0.010* |
|  | H2 | one-tail (↑) | 3.33% | 2.83% | 11 | 4.07 | 0.004* |
|  | H3a | one-tail (↑) | 4.84% | 2.93% | 11 | 5.71 | <0.001* |
|  | H3b | one-tail (↑) | 4.02% | 1.78% | 11 | 7.81 | <0.001* |
| Braking Asymmetry | H1 | one-tail (↓) | -12.06% | 5.47% | 11 | -7.63 | <0.001* |
|  | H2 | one-tail (↑) | 7.87% | 4.07% | 11 | 6.69 | <0.001* |
|  | H3a | one-tail (↑) | 17.24% | 7.54% | 11 | 7.92 | <0.001* |
|  | H3b | one-tail (↑) | 13.06% | 6.82% | 11 | 6.63 | <0.001* |
| Propulsion Asymmetry | H1 | one-tail (↑) | 2.57% | 3.13% | 11 | 2.84 | 0.032* |
|  | H2 | one-tail (↓) | -1.37% | 5.71% | 11 | -0.83 | 0.85 |
|  | H3a | one-tail (↓) | -5.29% | 3.06% | 11 | 6.00 | <0.001* |
|  | H3b | one-tail (↓) | -4.10% | 3.15% | 11 | -4.51 | 0.002* |

Paired Tests: H1 (ON:Early-OFF:Base), H2 (ON:Late-ON:Early), H3a (OFF:Early-ON:Late), H3b (OFF:Early-OFF:Base)
\* indicates statistical significance

*Hypothesis 1*, which stated that inducing asymmetry in gait kinematics and kinetics through the application of asymmetric hip stiffness with the exoskeleton is similar to the split-belt treadmill paradigm, was assessed using a paired *t*-test between the OFF:Base and ON:Early conditions for each measure.

*Hypothesis 2*, which stated that individuals would adapt their gait behavior back towards restoring gait symmetry in response to the application of asymmetric hip joint stiffnesses, was assessed using paired *t*-tests between the ON:Early and ON:Late conditions for each measure.

*Hypothesis 3*, which stated that individuals would exhibit an aftereffect in an opposite direction to initially induced asymmetry after the asymmetric hip joint stiffnesses is removed, was assessed using two paired *t*-tests for each measure. The comparison between the ON:Late and OFF:Early conditions tested whether there was a change in asymmetry in the opposite direcion to the initially induced asymmetry upon removal of the asymmetric hip joint stiffnesses (*Hypothesis H3a*), and the comparison between the OFF:Base and OFF:Early conditions tested whether the magnitude of aftereffect induced differed from the baseline level of symmetry as expected (*Hypothesis H3b*).

To control for Type I errors, a Bonferroni correction was applied to the reported p-values (referred to as *p*_*adjusted*_) for the four planned comparisons tests used. A custom MATLAB script was used to conduct all statistical analyses, and the significance level was set at *α* = 0.05 for all tests.

## III. RESULTS

### A. ANOVA Results

The ANOVAs revealed a statistically significant effect of condition on asymmetry for all kinematic and kinectic asymmetry measures, except for ankle ROM asymmetry. These results are summarized in Table I.

### B. Planned Comparisons

Fig. 2 illustrates how the kinematic and kinetic asymmetry measures changed over strides and across conditions for a representative subject. The results of the planned comparisons are summarized in Table II.

**Fig. 2.**
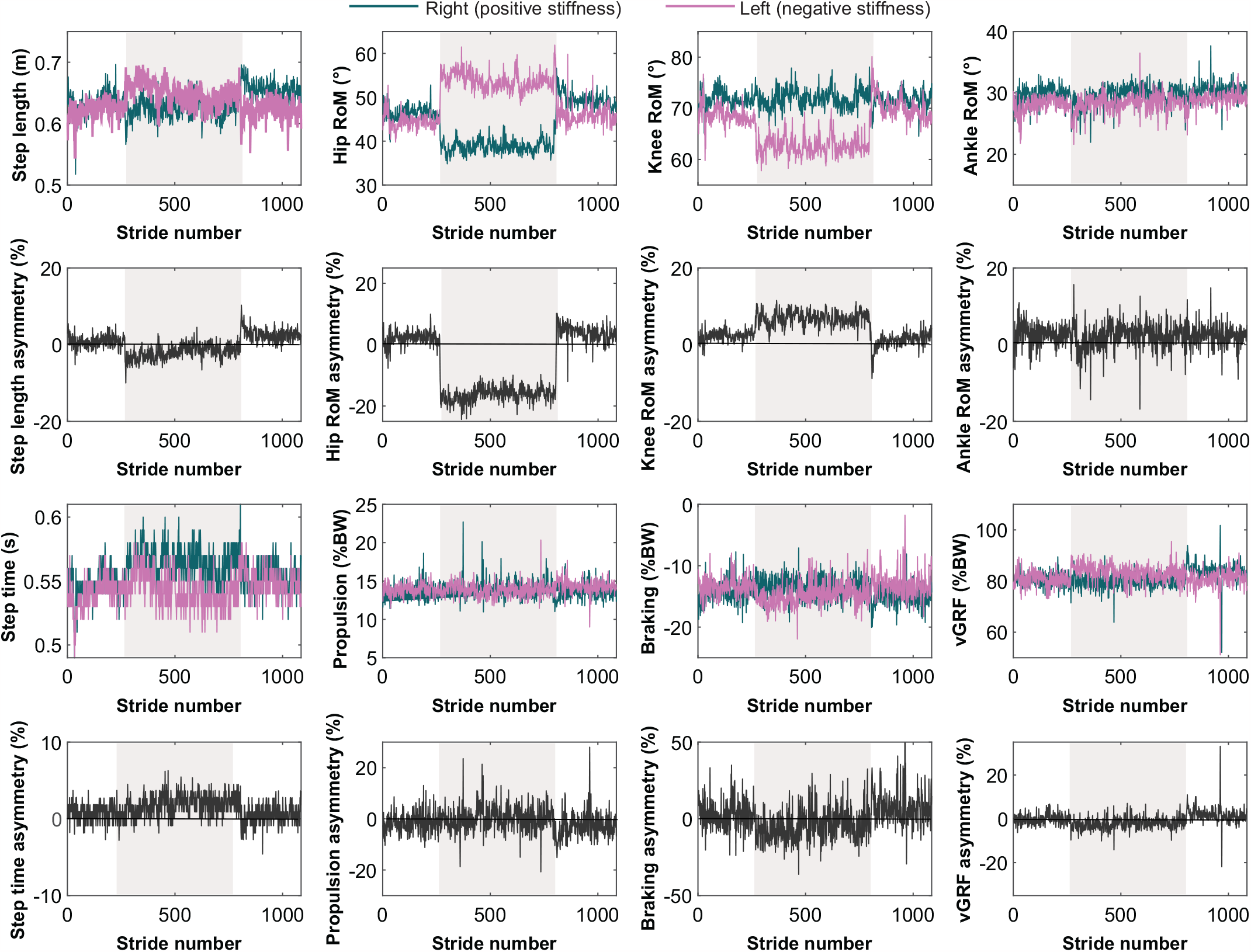
Dependent measures of a representative participant. Shaded regions represent when the stiffness controller was on.

*1)Hypothesis 1*

Consistent with *Hypothesis 1*, applying asymmetric stiffnesses about the hip joints induced asymmetries similar those induced by the split-belt treadmill paradigm. As predicted, the intervention induced a statistically significant negative asymmetry in step length and positive asymmetry in step time during the ON:Early condition compared to the OFF:Base condition (Figure 3). Step length was shorter and step time was higher for the positive stiffness leg compared to the negative stiffness leg. ^1^ A statistically significant negative asymmetry in hip ROM was induced as predicted, which was accompanied by a statistically significant positive asymmetry in knee ROM (Figure 4). Hip ROM was reduced and knee ROM was increased for the positive stiffness leg compared to the negative stiffness leg.

**Fig. 3.**
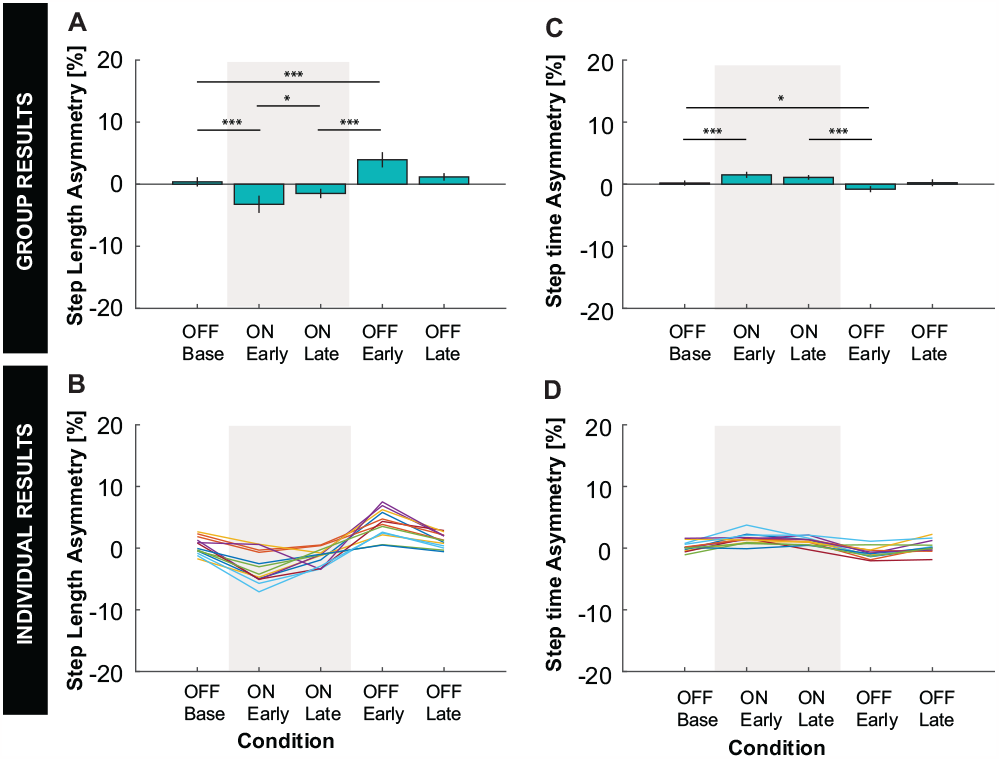
Step length and step time results. **A:** Group average and **B:** individual results for step length asymmetry. **C:** Group average and **D:** individual results for step time asymmetry. **A, C:** Error bars represent two standard errors of the mean. Shaded regions represent when the stiffness controller was on. The ANOVAs found a statistically significant effect of condition on all spatiotemporal measures. *, **, and *** indicate that the Bonferroni-corrected planned comparison between conditions was statistically significant with ***p***_***adjusted***_ ***<* 0.05, *p***_***adjusted***_ ***<* 0.01, *p***_***adjusted***_ ***<* 0.005**, respectively. **B, D:** Color indicates the different individual subjects.

**Fig. 4.**
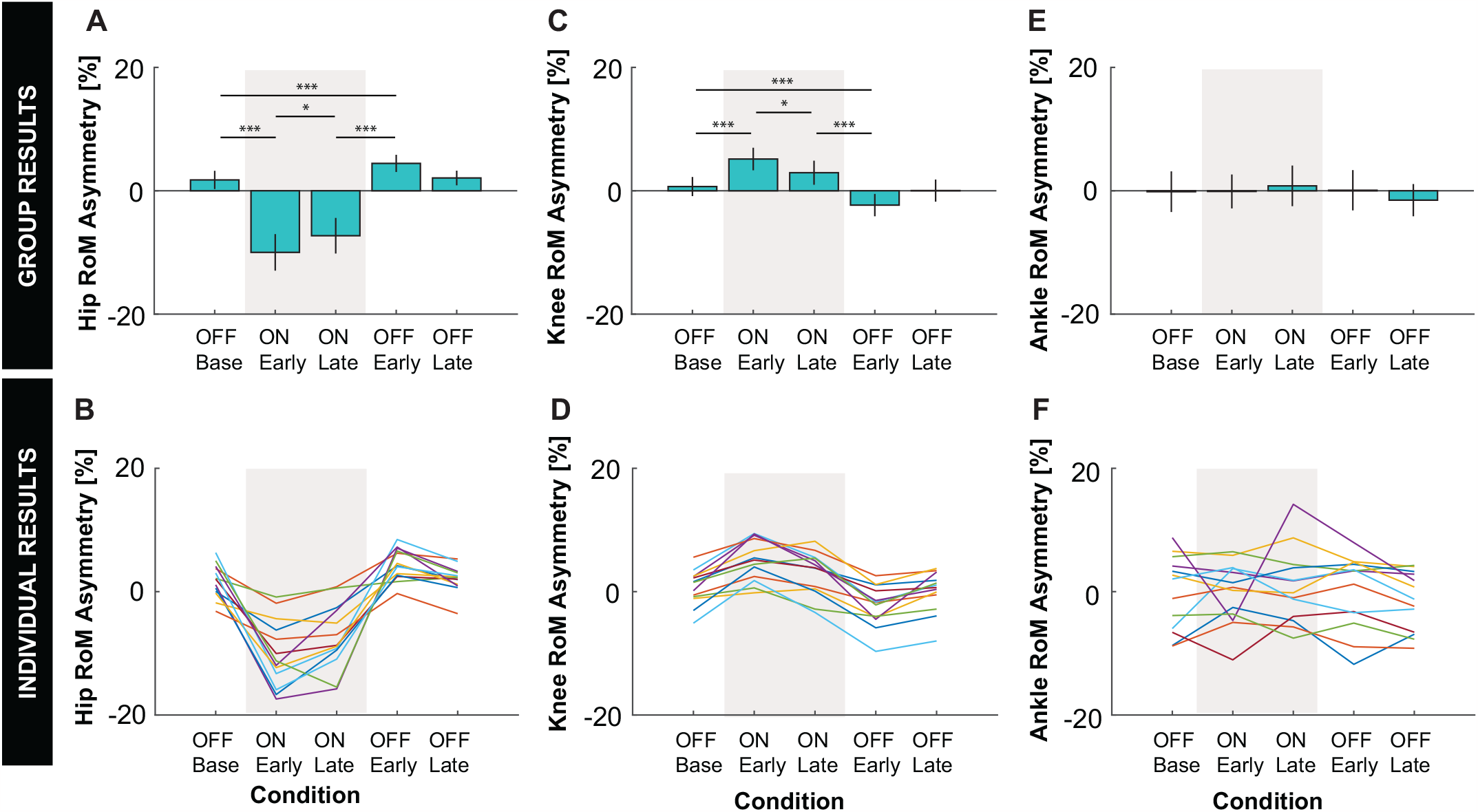
Joint kinematics results. **A:** Group average and **B:** individual results for hip RoM asymmetry. **C:** Group average and **D:** individual results for knee RoM asymmetry. **E:** Group average and **F:** individual results for ankle asymmetry. **A, C, E:** Error bars represent two standard errors of the mean. Shaded regions represent when the stiffness controller was on. The ANOVAs found a statistically significant effect of condition on all spatiotemporal measures. *, **, and *** indicate that the planned comparison between conditions was statistically significant with ***p***_***adjusted***_ ***<* 0.05, *p***_***adjusted***_ ***<* 0.01, *p***_***adjusted***_ ***<* 0.005**, respectively. **B, D:** Color indicates the different individual subjects.

Applying asymmetric hip stiffnesses also induced statistically significant negative asymmetries in vGRF and braking, as well as a statistically significant positive asymmetry in propulsion, compared to baseline as predicted (Figure 5). vGRF and braking forces were lower for the the positive stiffness leg compared to the negative stiffness leg, while the opposite was true for propulsive forces.

**Fig. 5.**
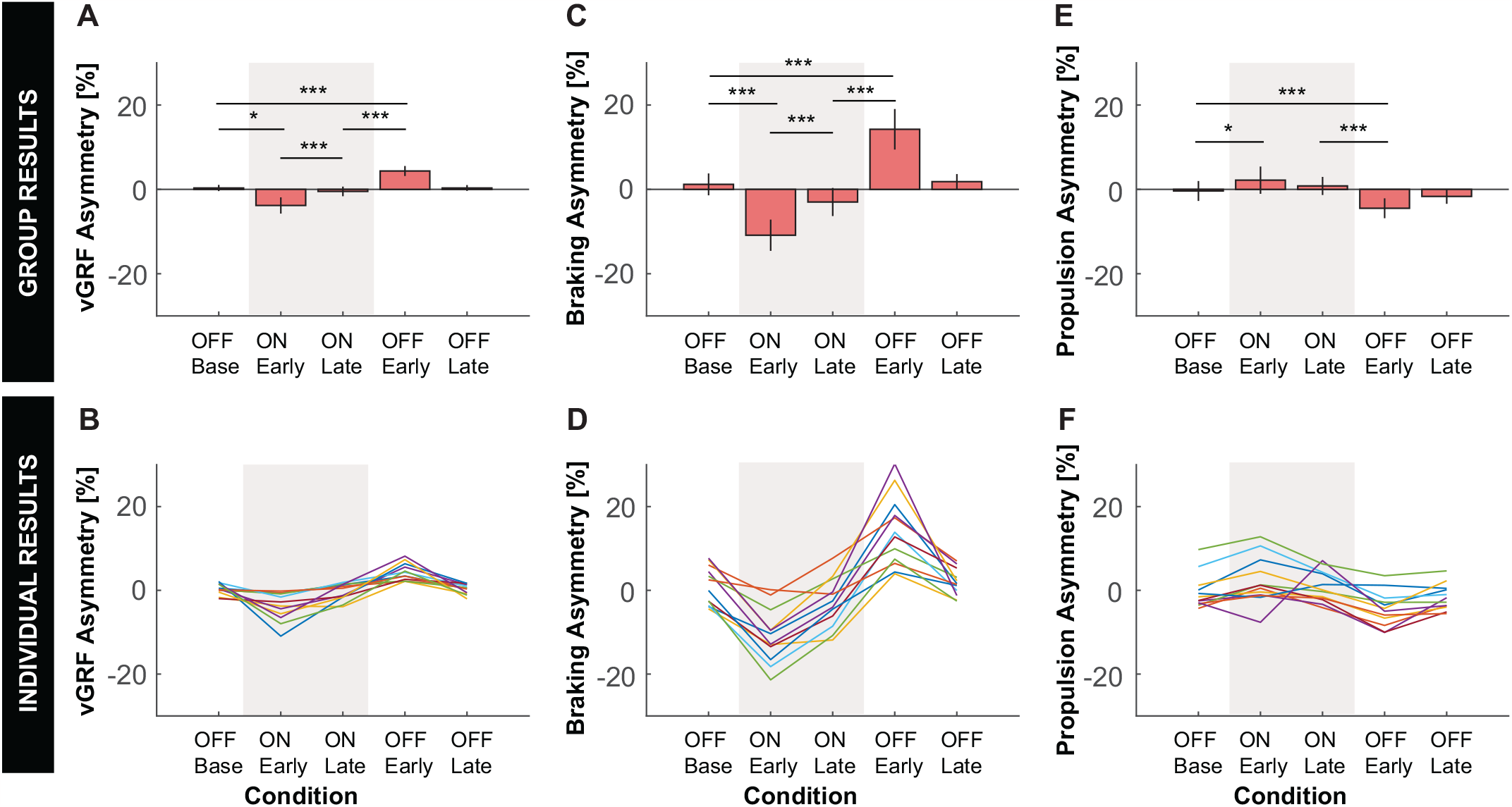
Kinetic results. **A:** Group average and **B:** individual results for propulsion asymmetry. **C:** Group average and **D:** individual results for braking asymmetry. **E:** Group average and **F:** individual results for vertical GRF (vGRF) asymmetry. **A, C, E:** Error bars represent two standard errors of the mean. Shaded regions represent when the stiffness controller was on. The ANOVAs found a statistically significant effect of condition on all spatiotemporal measures. *, **, and *** indicate that the planned comparison between conditions was statistically significant with ***p***_***adjusted***_ ***<* 0.05, *p***_***adjusted***_ ***<* 0.01, *p***_***adjusted***_ ***<* 0.005**, respectively. **B, D:** Color indicates the different individual subjects.

*2)Hypothesis 2*

Consistent with *Hypothesis 2*, participants responded to application of asymmetric hip joint stiffnesses by adapting their gait behavior back towards restoring symmetry. As predicted, statistically significant changes in the direction towards symmetry (i.e., 0% asymmetry) were observed in ON:Late compared ON:Early for all measures except step time and propulsion. While the mean changes in step time and propulsion were not statistically significant (*p*_*adjusted*_ = 0.200 and 0.845, respectively) and were small in magnitude, the mean changes were in the direction towards symmetry as predicted.

*3).Hypothesis 3*

Consistent with *Hypothesis 3*, aftereffects were observed for all measures after the stiffness controller was turned off. As predicted, there were statistically significant changes in asymmetry in the opposite direction to the initially induced asymmetry in the OFF:Early condition compared to the ON:Late condition (*Hypothesis 3a*). Moreover, the magnitude of aftereffect induced in the OFF:Early condition was significantly different from the baseline level of symmetry observed in the OFF:Base condition (*Hypothesis H3b*).

## IV. DISCUSSION

The goal of the study was to test whether the application of asymmetric hip stiffness would elicit signatures of neural adaptation similar to that of split-belt treadmill training [10], [26], [27], [29], [30]. As shown in Table II, our results illustrated that the intervention induced an immediate asymmetry in all kinetic and kinematic measures except for ankle RoM (Hypothesis 1). Over time, participants adapted towards a more symmetric gait pattern in most of the kinetic and kinematic measures (Hypothesis 2) (Table II). Turning off the stiffness controller induced an after-effect in an opposite direction to an initially exhibited asymmetry in all measures except for ankle RoM (Hypothesis 3). The induced after-effects washed out over time and participants changed their gait behavior back towards baseline levels of symmetry. By these criteria, our intervention successfully elicited the predicted response.

### A. Potential differences in adaptation compared to split-belt treadmill walking

Despite overall similarities, our results suggest potential differences in the adaptation to interlimb asymmetry induced at the hip joints compared to that induced at the feet. For instance, although participants adapted their gait towards symmetry in response to applied asymmetric hip stiffnesses, not every participant eventually reached baseline levels of asymmetry after 10 minutes of exposure.

One possible explanation is that complete adaptation to asymmetric stiffness at the hip joints requires more time compared to split-belt treadmill training. This could be attributed to the differences caused by torques applied directly at the hip joints versus speed constraints imposed on the feet during split-belt treadmill training. For instance, interactions with a split-belt treadmill are intermittent for each leg; foot motion is constrained by the moving belt during contact (stance phase), and unconstrained during non-contact (swing phase). In contrast, each leg maintains continuous contact with the exoskeleton. Moreover, the interaction between the user and each of these two devices differs. The exoskeleton will yield to the forces applied by the user, whereas the treadmill is only minimally responsive. Such differences impact the exploration possibilities afforded by each paradigm, potentially affecting the process of adaptation.

It is also possible that the nervous system adapts towards slightly different behaviors during each intervention. Such a difference could arise because (1) the objectives driving the adaptation differ during each intervention or (2) the same objective leads to distinct optimal behaviors for each intervention due to the differences described above.

Traditionally, split-belt treadmill adaptation has been described as an error (i.e., asymmetry) minimization process. However, recent studies have revealed that with longer adaptations periods (e.g., 45 minutes) individuals converge to behaviors beyond symmetry and begin walking with asymmetric step lengths in the opposite direction [31]. This behavior can be described as resulting from energy minimization, during which individuals learn to exploit the mechanical work performed by the treadmill to reduce their metabolic effort during walking [31]. It is also plausible that adaptation in split-belt treadmill walking is driven by a combination of asymmetry error reduction, metabolic cost minimization, and other objectives [32].

What objective(s) drive adaptation to applied asymmetric hip stiffnesses similarly remains an open question. It is possible that adaptation to asymmetric hip stiffnesses could be driven, at least in part, by energy minimization. Speculatively, participants may have maintained slightly asymmetric hip joint kinematics due to the added effort required to directly counteract motor torques. The observed behavior of convergence to asymmetric hip RoM and more symmetric step length enabled by compensations at the knee could therefore be interpreted as adaptation driven by a combination of step length symmetry and energy minimization. In contrast, recent results found no significant difference in hip, knee, or ankle RoM in the adapted gait behavior during split-belt treadmill walking compared to speed matched tied-belt walking in unimpaired individuals [33]. While this study demonstrates the nervous system’s ability to adapt to asymmetry induced by the hip exoskeleton, further investigation is required to elucidate the specific objectives driving adaptation, as well as the neural mechanism involved [34]. How these processes relate to those in split-belt treadmill walking and other gait asymmetry interventions will have important implications for potential gait rehabilitation.

### B. Limited adaptation of propulsion

Split-belt treadmill studies have shown a weak response in propulsion ground reaction forces. For instance, the presence of after-effects varied among different studies, with some showing them [26], [30] and others not [27], [29]. Similar to split-belt treadmill training, applying asymmetric stiffness at the hip joints also showed a weaker adaptation response in propulsion compared to the other measures. These findings underscore the importance of developing propulsion-based interventions, as individuals post-stroke often exhibit weight-bearing and propulsion asymmetries in addition to spatiotemporal asymmetries [4], [5].

### C. Promise of overground training to reduce gait asymmetry

The primary objective of rehabilitation is to achieve functional improvements that extend beyond the laboratory or clinical environment. Thus, developing interventions that improve gait symmetry during overground walking are critical. It is hypothesized that the limited transfer of symmetry improvements achieved with split-belt treadmill to overground walking is due to the difference in walking contexts [35]. Methods such as increased repetition [15] and increased cognitive load [36] have been shown to enhance the generalization of aftereffects from split-belt treadmill walking to overground walking. However, a more straightforward approach may be to train during overground, avoiding the need to transfer adapted behavior across walking contexts altogether.

Drawing direct inspiration from split-belt treadmill training, the development of asymmetry-inducing footwear has demonstrated the promise of overground training. For instance, walking with shoes that have asymmetric height has shown to induce asymmetries in temporal and lower extremity kinematic gait parameters [37]. Even though this study was performed on the treadmill, the results still show that this could be an effective overground training approach for individuals post-stroke. Also, overground training with passive footwear that generates a backward motion to the foot to exaggerate the step length asymmetry has been shown to improve step length asymmetry in individuals post-stroke [38]. A motorized version of such footwear has also been shown to induce adaptation in unimpaired individuals during treadmill walking [39]. As with split-belt treadmill walking, the potential clinical advantages and/or disadvantages of inducing asymmetry at the joint level compared to the endpoint (i.e., foot) during overground training remain to be determined.

In this study, we successfully demonstrated the feasibility of using a robotic hip exoskeleton to induce interlimb asymmetry and elicit neuromotor adaptation. While our experimentation took place on a dual-belt treadmill, the portability of our exoskeleton design opens the possibility of investigating overground walking beyond the confines of a laboratory setting in future studies. Consequently, future research will focus on evaluating the potential for longer persistence of adapted gait behavior during overground training with the robotic hip exoskeleton, as well as exploring its effectiveness as a gait rehabilitation tool.

## V. CONCLUSIONS

This study aimed to assess the impact of applying bilateral asymmetric stiffness using a hip exoskeleton on spatiotemporal and kinetic gait parameters among healthy individuals. The findings revealed behavioral signatures of neural adaptation, similar to those observed in split-belt treadmill training. Consistent with split-belt treadmill research, turning on the stiffness controller initially induced an immediate asymmetry across gait measures, except for ankle RoM. Over time, participants adapted towards a more symmetric gait pattern in most of the kinematic and kinetic gait measures. Turning off the stiffness controller elicited an after-effect in the opposite direction to an initially induced asymmetry, which washed out over time. Even though this study was conducted on a dual-belt treadmill, it still showed meaningful results about how humans adapt to asymmetric stiffness applied by a hip exoskeleton. Forthcoming research will focus on investigating the effectiveness of proposed intervention during overground walking, outside of the confines of a laboratory setting.

## ACKNOWLEDGMENT

We would also like to thank Dasha Trosteanetchi for her assistance in processing motion capture data for this study.

## Footnotes

1 At first glance, this might seem counterintuitive because one might anticipate step length and step time to change together. However, it is important to recognize that this discrepancy arises from the conventional definition of step time in the field. We continue to use this measure for the sake of consistency and to facilitate comparisons with existing literature.

